# Supervised machine learning with feature selection for prioritization of targets related to time-based cellular dysfunction in aging

**DOI:** 10.1101/2022.06.24.497511

**Authors:** Nina Truter, Zuné Jansen van Rensburg, Radouane Oudrhiri, Raminderpal Singh, Carla Louw

**Author notes:** Nina Truter and Zune Jansen van Rensburg these authors contributed equally to this work.

## Abstract

**Background:** Global life expectancy has been increasing without a corresponding increase in health span and with greater risk for aging-associated diseases such as Alzheimer’s disease (AD). An urgent need to delay the onset of aging-associated diseases has arisen and a dramatic increase in the number of potential molecular targets has led to the challenge of prioritizing targets to promote successful aging. Here, we developed a pipeline to prioritize aging-related genes which integrates the plethora of publicly available genomic, transcriptomic, proteomic and morphological data of *C. elegans* by applying a supervised machine learning approach. Additionally, a unique biological post-processing analysis of the computational output was performed to better reveal the prioritized gene’s function within the context of pathways and processes involved in aging across the lifespan of *C. elegans*.

**Results:** Four known aging-related genes — daf-2, involved in insulin signaling; let-363 and rsks-1, involved in mTOR signaling; age-1, involved in PI3 kinase signaling — were present in the top 10% of 4380 ranked genes related to different markers of cellular dysfunction, validating the computational output. Further, our ranked output showed that 91% of the top 438 ranked genes consisted of known genes on GenAge, while the remaining genes had thus far not yet been associated with aging-related processes.

**Conclusion:** These ranked genes can be translated to known human orthologs potentially uncovering previously unknown information about the basic aging processes in humans. These genes (and their downstream pathways) could also serve as targets against aging-related diseases, such as AD.

## Introduction

The global life expectancy has increased by more than 6 years in the last two decades, without a corresponding increase in health span (life without major disease or disability) (1). With an expected near doubling in the number of people over the age of 60 years globally by 2050, this poses a major socioeconomic burden and an urgent need exists to delay the onset of age-related diseases such as Alzheimer’s disease (AD) and cardiovascular disease (1).

A major drive for the implementation of interventions that increase health span and delay senescent span is therefore underway (Figure 1; modified from (2) for *C. elegans*), with the aim to promote successful and to limit unsuccessful aging. Successful aging can be defined as the decline in cellular, tissue, and organ function over an organism’s lifespan without the onset of pathology and with the presence of high physical, cognitive, and social function (adapted from (3) using (4)). An improved understanding of the cellular and molecular mechanisms and the rate of their deterioration is needed to develop therapies for successful aging. This requirement has contributed to the definition of the nine hallmarks of aging, which include intracellular processes that manifest during normal aging (5). These include: genomic instability, telomere attribution, epigenetic alterations, loss of proteostasis, deregulated nutrient-sensing, mitochondrial dysfunction, cellular senescence, stem cell exhaustion, and altered intercellular communication (5).

**Figure 1:**
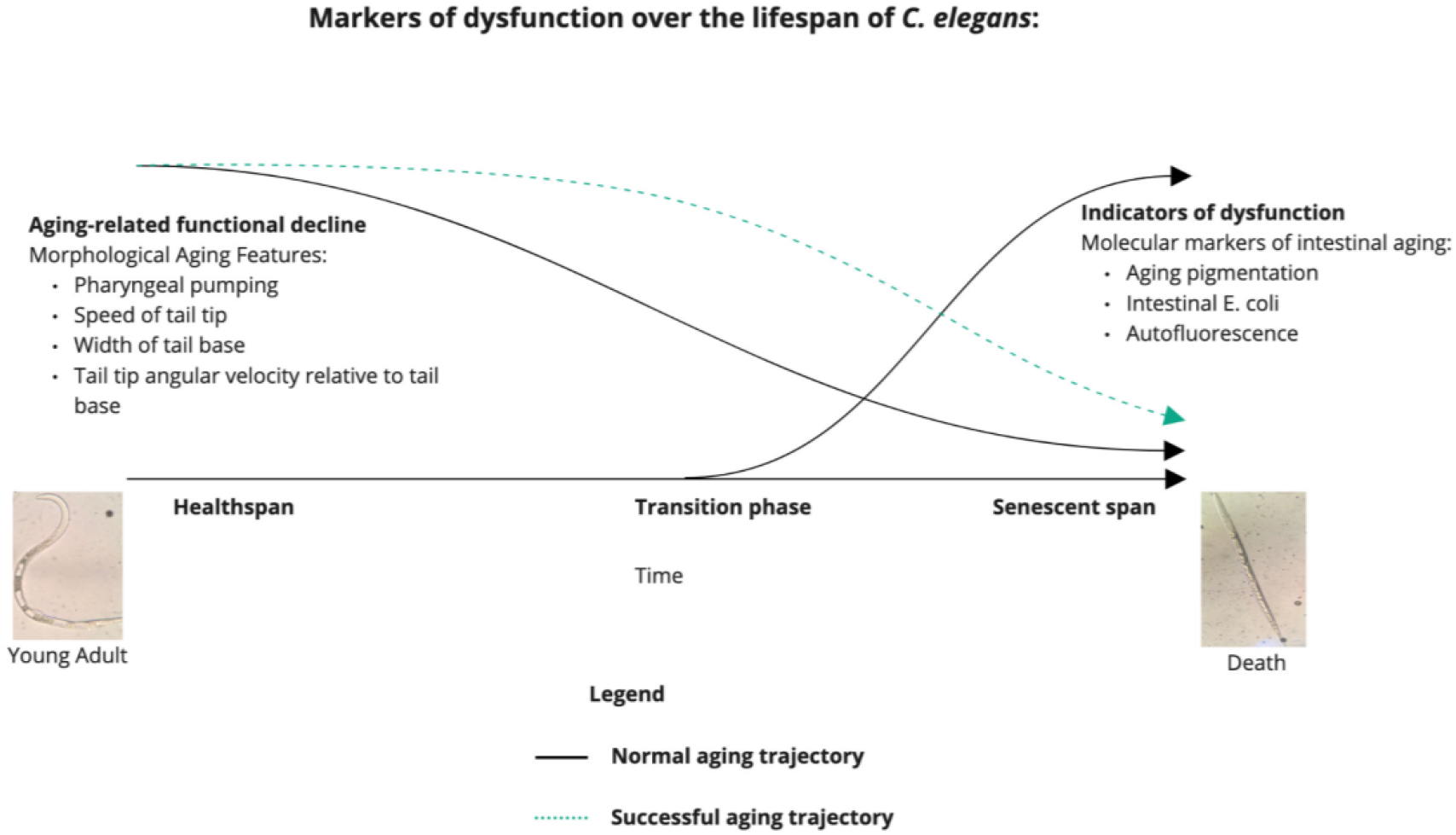
Markers of dysfunction observed over the lifespan of C. elegans related to the aging process, specific to this study. Aging-related functional decline includes morphological aging features which degrade over C. elegans’s lifespan and other indicators of dysfunction which include molecular markers of intestinal aging that accumulate over C. elegans’s lifespan. The black line indicates the normal aging trajectory in C. elegans, while the green dotted line indicates a successful aging trajectory in C. elegans.

Since the introduction of the above hallmarks of aging, potential molecules which could modulate the activity of mechanistic pathways involved in aging have been reported. The GenAge database summarizes targets associated with human longevity and currently describes 307 genes (6). These targets include genes from known age-associated pathways, such as the sirtuin, insulin/IGF-1 signaling, AMP-activated protein kinase (AMPK), and mTOR pathways (7,8). Although some success has been achieved by targeting components in these pathways with promising results in clinical trials, such as inhibiting mTOR by rapamycin (9), the relative contribution of each hallmark and consequently its involved molecular mechanisms driving and contributing to the overall aging process remains unclear (10). This makes the prioritization of targets for therapeutic intervention challenging. Moreover, longitudinal studies of human aging as well as the development of pharmacotherapies are time consuming and costly (11). Given these challenges, a particular need arises to better prioritize targets based on their ability to modulate the aging process, without solely relying on *in vitro* and *in vivo* experiments (11,12).

To address these challenges, increased utilization of database searches, biological assays, and machine learning techniques have been implemented to better identify targets of interest (12–15). These techniques have been used to advance research in the field of neurodegeneration, with a specific focus on AD. Targets in AD have been identified through a multi-omics approach focusing on protein networks and pathways during disease progression (15). Semi-supervised machine learning has also been used to identify an MRI biomarker for mild cognitive impairment (MCI), effectively predicting the clinical conversion from MCI to AD up to three years prior to disease onset (16). MRI imaging data has also been used in an ML approach to aid in disease diagnosis (17). Given the complexity of the biological systems and the wealth of available data, the use of machine learning seems invaluable to accurately and efficiently identify targets for ND.

The careful consideration of targets identified with machine learning techniques in the context of biological processes is necessary to assess the feasibility and actionability of the identified targets. One such approach is taken by Pun *et al*., (2022) by using the PandaOmics platform to investigate targets associated with aging and age-associated diseases from human cross-sectional data based on the nine hallmarks of aging (12). Although the use of human data to identify targets is highly relevant for drug discovery, obtaining rich longitudinal data to understand disease progression across an individual’s lifespan has remained extremely challenging. Current approaches are therefore typically reliant on cross-sectional data, further limiting the deduction of cellular dynamics in aging. Short-lived simple animal models can, however, aid in circumventing this limitation as they are ideal for studying the intracellular evolutionarily conserved processes of aging (18,19).

*C. elegans* is one of the most popular simple organisms used to investigate aging (8). Its short lifespan enables the in-depth characterization of its life stages to generate time-based data on different levels of Omics (20,21). Importantly, several of the intracellular hallmarks of aging are conserved across species. Despite *C. elegans* lacking some of the evolutionary advanced aging hallmarks present in other multicellular organisms due to the postmitotic nature of many of its cells, the core hallmarks of aging can be investigated in a multicellular organism with less complexity than found in mammalian species (18). As the hallmarks of aging are interconnected, many of the non-conserved, more advanced hallmarks exert their eventual effects through conserved pathways (18). Therefore, understanding the core mechanisms of aging in isolation can inform strategies that aim to improve their function, in order to promote successful aging (Figure 1). Altered insulin signaling by knockout of daf-2, for example, resulted in a reduced accumulation in aging pigment (lipofuscin) and improved locomotor capacity, which is indicative of successful aging (22).

A further benefit of working with *C. elegans* is the vast range of available data including time-based transcriptomic, proteomic, phosphorylation status, molecular data and detailed morphological data, allowing to describe the relationships between different Omics layers and their possible contributions to the aging process (20,21,23,24). However, prioritization of aging-related targets remains a challenge in this organism due to the many existing targets and their association with longevity and not necessarily the aging process.

In this study, supervised machine learning and recursive feature elimination techniques were used to construct a pipeline to prioritize genes associated with aging and thereby identify potential targets. To achieve this, several time-based biological layers, including markers of cellular and macroscopic function were used to characterize the aging process in *C. elegans*. This included a broad range of categorical and time-dependent phenotypic and genotypic data; i.e. RNA and protein levels, phosphorylation status, and the biological process description of each gene. A unique post-processing workflow was established to translate the prioritized genes to pathways and cellular processes in human aging, to support their potential druggability for aging-related diseases.

## Methods

A computational pipeline (workflow) was constructed which includes supervised machine learning on time-based data of *C. elegans* genes (feature variable) and aging-related genes (response variable) (Figure 2). An example of the computational output with post-processing using biological analysis is presented below.

**Figure 2:**
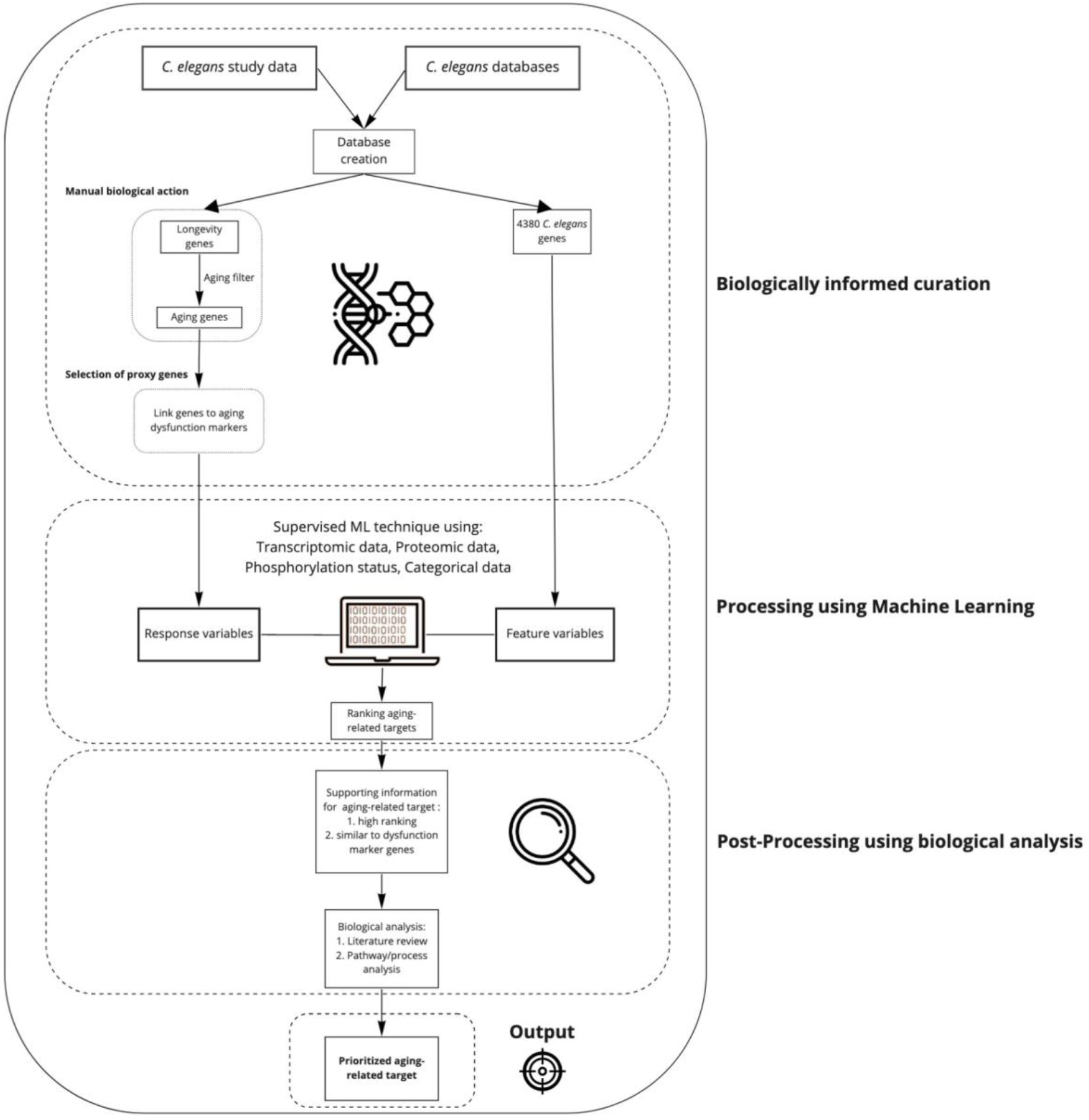
Workflow demonstrating the synergy between biological analysis and supervised machine learning using time-based C. elegans data to prioritize aging-related targets using: i. Biologically-informed curation of data from C. elegans studies and databases, used to select genes for analysis. ii. Processing of aging-related and proxy genes (response variables) and 4380 general genes (feature variables) to produce a ranking of aging-related targets. iii. Post-processing of aging-related genes to include supporting information for their use as aging-related targets through biological analysis to produce a prioritized aging-related target.

### Data collection and formatting

Data were collected from open-source databases and from literature. The data collected from different studies (Table 1) were combined with the Wormbase Gene IDs or alternatively the gene names. Several layers of Omics were used to understand the broader context in which the targets could exert effects on the aging process (Figure 3). All computational analyses were conducted in Python 3.8 (25). MinMaxScaler from the Scikit-learn package was used to standardize the combined data (26). Time course data were used in the original format while the categorical data were converted to binomial data by One-hot encoding (26). Pandas and Numpy packages were used for data formatting (27,28). Genes with missing data fields were excluded from the dataset. A total number of 4380 active *C. elegans* genes were further included in computational analyses (Figure 2).

**Table 1:** Datasets used for supervised machine learning model training and their sources. “Categorical” data refers to quantitative data converted to binomial data and “numerical” data refers to time course datasets.

| Datafield | Source |
| --- | --- |
| Gene IDs and Gene names | Wormbase (29) |
| Biotype (categorical) | Wormbase (29) |
| Expression pattern tissue (categorical) | Wormbase (29) |
| Expression pattern lifestage (categorical) | Wormbase (29) |
| Expression pattern lifestage (numerical) | Wormbase (29) |
| Gene ontology association (categorical) | Gene ontology (30,31) |
| Morphological feature change over time (numerical) | Martineau (23,32) |
| Proteome ratios over time (numerical) | Narayan (20) |
| Gene expression over time (numerical) | Hastings (21) |
| Gene expression over time (numerical) | Bgee (33) |
| Gene effect on lifespan (categorical) | GenAge (6,34) |
| Protein Phosphorylation (numerical) | Huang (24) |
| Pathway labels (categorical) | Uniprot (35) |
| Interacting genes (categorical) | Wormbase (29) |
| Disease information (categorical) | Wormbase (29) |
| Human ortholog (categorical) | Wormbase (29) |
| Gene description (categorical) | Wormbase (29) |

**Figure 3:**
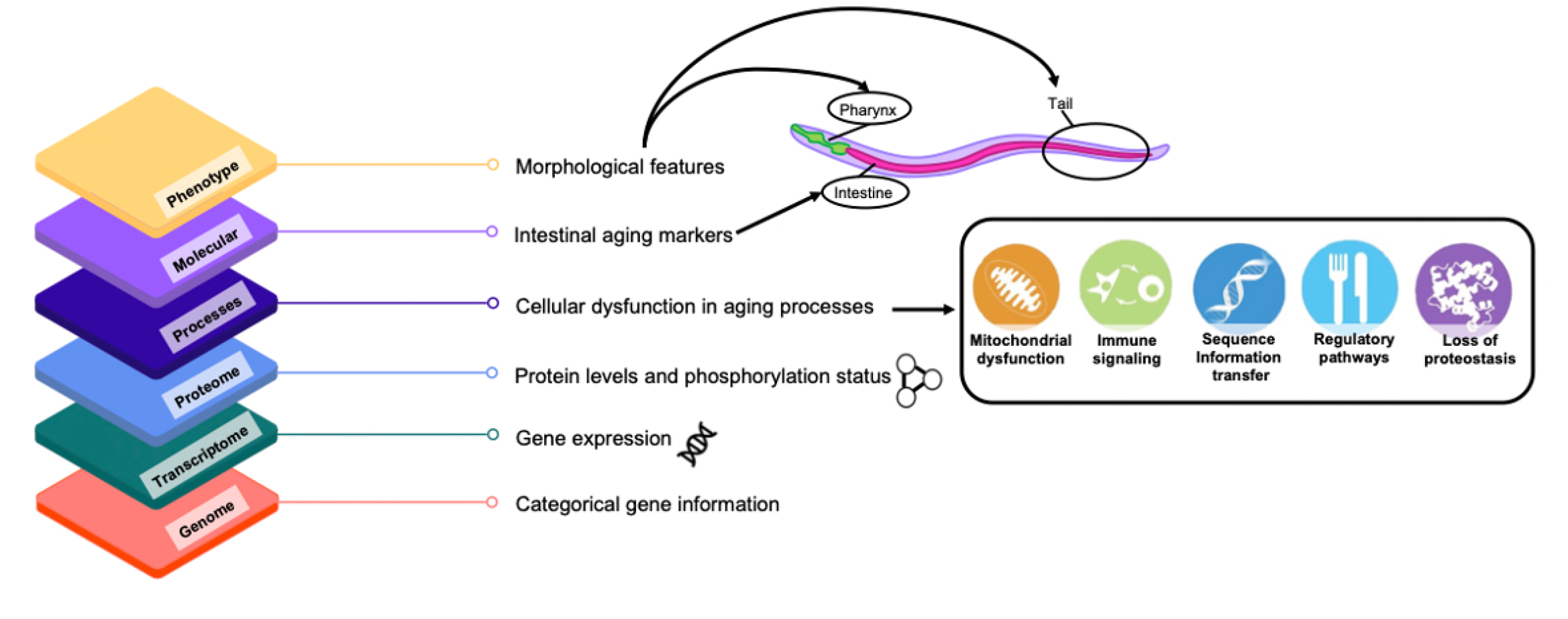
Visual representation of the different Omics layers used in the presented pipeline and the data used to characterize them.

### Compiling a concise list of genes associated with processes of aging

A list of genes associated specifically with aging processes was required as response variables for training of a supervised machine learning model to highlight targets within aging processes (Figure 2). First, a complete dataset of the known life-extending genes (n=887) was utilized from the public database GenAge (6), https://genomics.senescence.info/genes/stats.php, accessed Feb 2022. Only 429 genes had the required data (categorical information, gene expression, protein levels and phosphorylation status) to be used within our computational model. However, due to the discrepancy between aging and longevity (36) the life-extending (longevity) genes were manually filtered to compile a list of genes consisting only of genes associated with known aging processes. Genes were chosen if their function listed on UniProt or Wormbase formed part of a known intracellular process associated with the hallmarks of aging (5,18,35). These processes of aging included: mitochondrial dysfunction (mitochondrial integrity and biogenesis and reactive oxygen species), sequence information transfer (a combination of the hallmarks genome instability and epigenetic alterations), immune signaling (the intracellular component of inflammation), loss of proteostasis and regulatory pathways (including deregulated nutrient sensing) (Figure 3). Once completed, the list contained 378 genes that function within a known aging process (Supplementary Table 1). These genes were used as response variables during model training, either as proxy genes associated with dysfunction markers or as non-specific aging associated genes.

### Dysfunction markers representative of aging

Various markers of aging indicative of organism dysfunction in *C. elegans* over time were used to understand how genes could contribute to the aging process, not only on a genetic level but also on a physiological/functional basis. Three types of dysfunction markers were selected across omics layers: either macroscopic morphological markers (“morphological aging features”), molecular markers (“intestinal aging”) or genetic markers (“genes associated with aging processes”) (Figure 3).

#### Morphological features

Morphological features are used to quantify the macroscopic health of *C. elegans* over time as it ages (32). Furthermore, changes in morphological features have been shown to be related to cellular function, for example, changes in dopaminergic neuronal function in knock-out studies are evident in locomotion (37). Both locomotion and pharyngeal pumping are related to lifespan and possibly to each other, with both showing a reduction in speed over time (38). Pharyngeal pumping and three features related to locomotion that decline significantly during aging (speed of tail tip, width of tail base and tail tip angular velocity relative to tail base) were included as dysfunction markers (32,39).

#### Markers of intestinal aging

The *C. elegans* intestine is the location of many stress responses that change during aging (40). Several known *C. elegans* aging markers are associated with the intestine, including autofluorescence and aging pigment (22,41). Furthermore, the accumulation of *E. coli* in the intestinal lumen is related to the lifespan of *C. elegans (42)*. An increase in autofluorescence, aging pigment or *E. coli* accumulation are indicative of general cellular dysfunction during aging, including a decline in immune signaling and redox balance dysfunction (22,41,42).

#### Non-specific genes associated with the aging process

Genes from the list of 378 aging-related genes that did not form part of the morphological or intestinal aging markers were used as “genes associated with aging processes” markers (n=329).

### Selection of proxy genes for dysfunction markers

Although all 378 aging-associated genes were used for model training, we wished to interpret the computational output within the context of dysfunction markers associated with aging. Therefore, seven proxy genes were selected for the markers of dysfunction based on computational analysis. Next, in order to identify proxy genes computationally, we assumed that a change in a dysfunction marker over time will be highly correlated (positively or negatively) with the change in expression over time of the genes associated with the respective marker. Based on this assumption we selected the genes with the lowest (closest to zero) Root Mean Square Error (RMSE) between the change in gene expression and dysfunction marker over time. Only seven proxy genes could be found with RMSE values < 0.3 for some dysfunction markers, therefore seven genes were selected for each marker to ensure equal contributions towards the ranking process by each dysfunction marker. Gaussian Process Regression was used to infer gaps in temporal data (21,22,32,39,41,42) and Linear regression was used to calculate the RMSE values.

Importantly, to ensure that the genetic association with the morphological aging features were biologically sensible, the genes selected to represent the feature had to be involved in the respective process or expressed within the corresponding tissue (based on Wormbase information and (43). Specifically, genes with a high RMSE value with involvement in movement and/or with expression in the tail were used for the features “speed of tail tip”, “width of tail base”, “tail tip angular velocity relative to tail base”. Similarly, genes expressed in the pharynx were used for pharyngeal pumping (Wormbase information and (43).

### Gene ranking with recursive feature elimination

All genes were ranked according to their similarity to the dysfunction marker genes. This similarity was determined by Recursive Feature Elimination (RFE), a feature selection algorithm. During RFE a machine learning model was iteratively retrained while the weakest predictive “feature” (in this case gene) was removed with each iteration. This process aids in the training of cleaner, more effective models, eliminating unnecessary features. RFE outputs a list of all features (genes), ranked from most relevant to least relevant for the model’s predictive capability. In this study, RFE was used to eliminate genes that contribute least to the processes of aging-associated dysfunction, during model training. Instead of using the traditional elimination of unnecessary features, we eliminated unnecessary genes, generating an output of genes ranked from most similar to the dysfunctions markers to least similar. A Support Vector Machine and Generalised Linear models were used within the RFE wrapper algorithm (26).

### Validation of genes associated with aging processes ranking

Genes from the GenAge database known to be associated with processes of aging were ranked, along with all other genes, by their predicted association with aging processes. The model was validated by its ability to highly rank genes known to be associated with aging. This validation is visualized in Figure 4 - the predicted ranking of these genes are distributed across 10 quantiles. We defined accurate predictive capability of the model if >75% of known genes associated with aging processes were ranked in the top three quantiles (30%) given the ranking output of the analyses. To further validate the computational ranking output, the top three genes from GenAge (modulating their expression results in the largest life extension) were selected: daf-2 (insulin receptor-like gene), age-1 (PI3 kinase signaling), and let-363 (mTOR signaling). The positions of these genes within our ranked output were determined.

**Figure 4:**
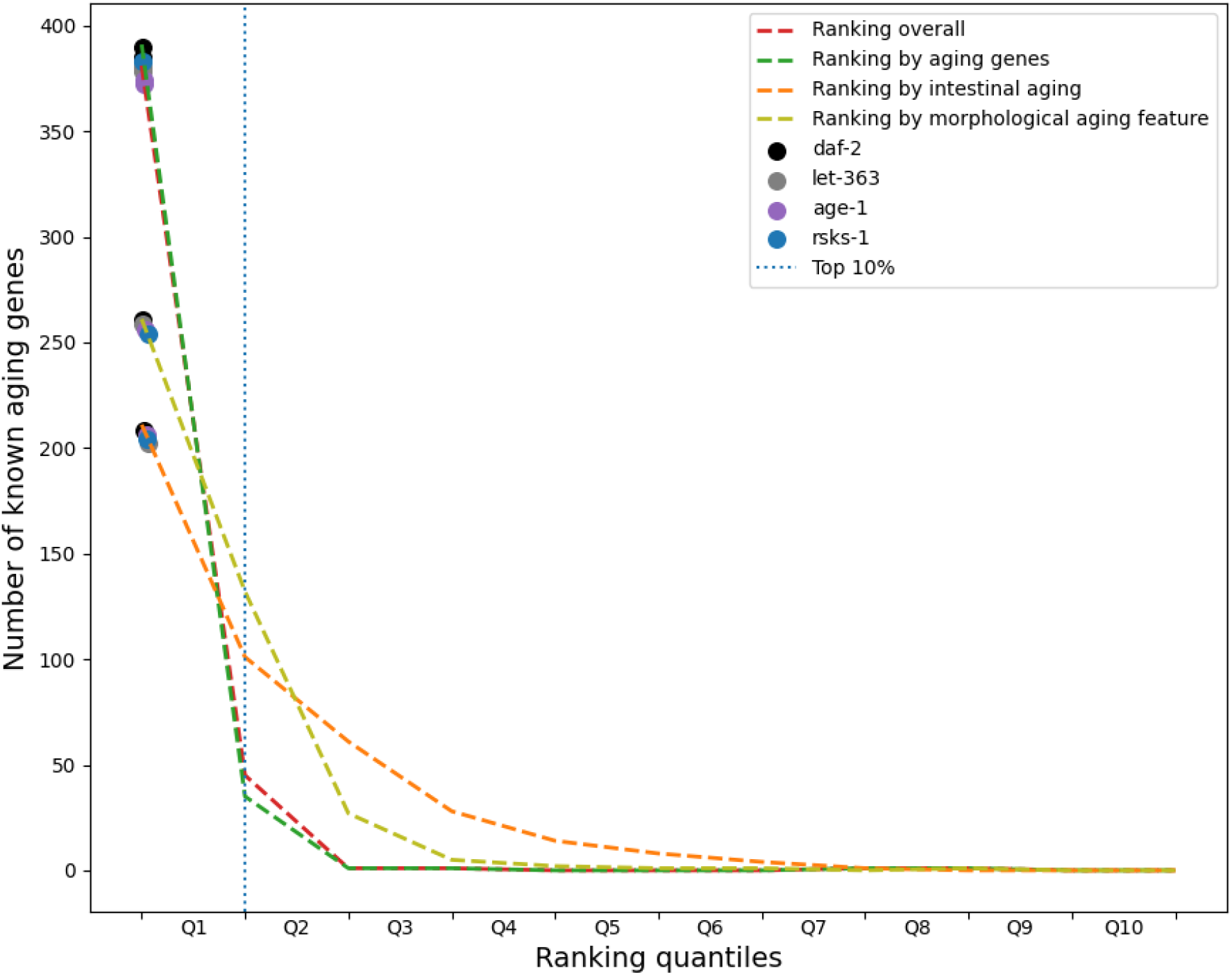
Ranking of known genes associated with aging processes (6,34) by the model as validation of the ranked output. The number of known aging-related genes placed in each of the 10 quantiles of the ranked output is shown by different dysfunction marker types: (aging-related genes (green), intestinal aging (orange), and morphological aging (light green) feature) and across all three dysfunction marker types (overall; red). The rankings of the three known aging-related genes, daf-2 (black), let-363 (gray), and age-1 (purple), as well as the selected aging-related gene, rsks-1 (blue), are shown by each dysfunction marker type and overall.

### Rsks-1 as example aging target

The computational output includes an overall ranking of the gene’s strength of association with known genes from aging processes. Furthermore, the similarity of the ranked genes to specific dysfunction marker genes is included. The output also includes correlation analyses between the chosen gene’s mRNA expression, protein levels, phosphorylation status, and categorical information, and those of all associated dysfunction markers. This can potentially associate the ranked gene more strongly with a dysfunctional process based on the strength of the similarity. In order to validate the computational output, a known gene associated with the aging process of the mTOR pathway, rsks-1, was selected. This selection was made since rsks-1 was ranked in the top 100 genes of the computational output, had been reported in relatively few publications (16 publications on Pubmed, https://pubmed.ncbi.nlm.nih.gov/, in the last 5 years), and its human ortholog has existing drugs targeting its activity (S6K1 inhibitors) (44,45).

### Discovery of known and predicted associations between genes (matrix)

Possible associations between the chosen gene (rsks-1) and dysfunction marker genes with the highest similarity were explored. This was done by placing the dysfunction marker genes and rsks-1 into their designated aging processes used for our list of aging-related genes (Supplementary Table 1). The aging processes attributed to the genes were chosen based on its Uniprot and Wormbase biological description (35). The known association between genes (Pubmed) or predicted associations based on computational output were visualized in a diagram for further interpretation.

In summary, a supervised machine learning model was trained to rank genes based on their similarity to known aging-related genes from GenAge. These known aging-related genes are grouped into dysfunction marker categories: morphological features, intestinal aging, and non-specific aging processes. Genes can be ranked by their similarity to all known aging genes or by the sub-groups of genes in the dysfunction marker categories.

## Results

### Assigning genes to dysfunction marker categories

Dysfunction markers of aging were selected in three categories based on their biological relevance to aging. For the categories (i) Intestinal aging and (ii) morphological aging feature, their respective seven proxy genes are listed in Table 2. All other genes formed part of the (iii) genes associated with aging processes category.

**Table 2:** Dysfunction marker types and their respective markers representative of aging. Dysfunction marker proxy genes associated with the eight dysfunction markers were selected from literature and regression analyses [Ref: Methodology_Selection of proxy genes for dysfunction markers]. The RMSE values of the regression analyses are included in brackets.

| Dysfunction marker type | Dysfunction marker | Dysfunction marker proxy genes (RMSE values) |
| --- | --- | --- |
| (i) Intestinal Aging | Aging pigmentation | age-1 (0.15), atg-18 (0.17), atg-9 (0.18), daf-2 (0.14), hcf-1 (0.22), hlh-30 (0.17), unc-51 (0.07) |
|  | Intestinal *E. coli* | akt-1 (0.23), daf-8 (0.19), dcr-1 (0.24), ire-1 (0.25), lgg-1 (0.13), sip-1 (0.18), wwp-1 (0.21) |
|  | Autofluorescence | daf-15 (0.13), hmg-4 (0.09), hsf-1 (0.12), let-765 (0.11), lin-15B (0.11), mpk-1 (0.14), smk-1 (0.12) |
| (ii) Morphological Aging Feature | Pharyngeal pumping | cua-1 (0.26), elc-1 (0.22), fbp-1 (0.20), mrp-5 (0.25), mtch-1 (0.17), tmem-135 (0.28), vha-6 (0.23) |
|  | Speed of tail tip | arr-1 (0.13), bar-1 (0.11), blmp-1 (0.06), itr-1 (0.09), lys-8 (0.12), pat-10 (0.12), spl-1 (0.03) |
|  | Width of tail base | atp-2 (0.08), ced-3 (0.06), egl-30 (0.04), eat-6 (0.06), chn-1 (0.10), pat-4 (0.07), unc-64 (0.10) |
|  | Tail tip angular velocity relative to tail base | egl-27 (0.17), let-502 (0.18), apy-1 (0.09), glp-1 (0.19), npp-3 (0.16), daf-7 (0.18), unc-32 (0.09) |
| (iii) Genes associated with aging processes | Uncategorised Aging-related Genes* | Remaining 329 aging-related genes not included in another dysfunction marker classification. |

\*Uncategorised aging-related genes were selected from literature only, not by regression analysis.

### Ranking of *C. elegans* genes based on dysfunction marker proxy genes

Genes were ranked based on their similarity to aging-related dysfunction markers. The ranking of genes by their potential involvement in the process of aging (i. e. their potential importance in determining the appearance of the dysfunction markers) is shown in Supplementary Information (Supplementary Table 2).

### Validation of ranked output using known aging genes

The accuracy of the ranking of genes by their potential involvement in the aging process is validated by the model’s ranking of known aging-related genes. The ranking of known genes associated with aging processes from the GenAge database (6,34) is shown in Table 3. The majority of the known aging-related genes were ranked in the first three quantiles (n=1314) of the ranking output (Figure 4).

**Table 3:** Ranking of known aging-related genes (daf-2, let-363, and age-1), and the selected aging-related gene, rsks-1, by the model out of 4380 C. elegans genes. The ranks of the genes per dysfunction marker type are shown, as well as the ranking percentiles in brackets.

| Gene | Overall rank<br>[n out of 4380, %] | Morphological aging feature rank<br>[n out of 4380, %] | Intestinal aging rank<br>[n out of 4380, %] | Genes associated with aging processes rank<br>[n out of 4380, %] |
| --- | --- | --- | --- | --- |
| Known genes from aging processes |  |  |  |  |
| *daf-2* | 37 (1%) | 50 (2%) | 98 (3%) | 45 (2%) |
| *let-363* | 16 (1%) | 52 (2%) | 284 (7%) | 14 (1%) |
| *age-1* | 97 (3%) | 146 (4%) | 216 (5%) | 98 (3%) |
| Selected aging-related gene |  |  |  |  |
| *rsks-1* | 55 (2%) | 259 (6%) | 221 (6%) | 39 (1%) |

Three known aging-related genes (daf-2, insulin signaling; let-363, mTOR signaling; age-1, PI3 kinase signaling) and one general aging-related gene (rsks-1) were selected for further validation of the output. The rankings of these four genes are also shown (Figure 4) indicating their positions based on each dysfunction marker type (aging-related genes, intestinal aging, and morphological aging feature). All four genes were ranked in Q1 based on the overall ranking (mean of all rankings by the individual dysfunction marker genes) as well as the ranking by aging-related genes, morphological aging features, and intestinal aging (with a ranking of <400). The ranking of known aging-related genes out of the 4380 genes that were analyzed are shown in Table 3.

Figure 5 shows that the top quantile of ranked genes (Q1 in Figure 4) consists of genes that are listed on GenAge, known to be aging-related genes (91% of the Q1 genes) (6,34). The remaining 9% of the Q1 genes are, to the best of our knowledge, not yet associated with aging-related processes. Based on the gene descriptions of Uniprot, these genes are involved in a variety of functional processes that could potentially affect aging (35). Some of these processes include iron metabolism, protein degradation and amino acid metabolism.

**Figure 5:**
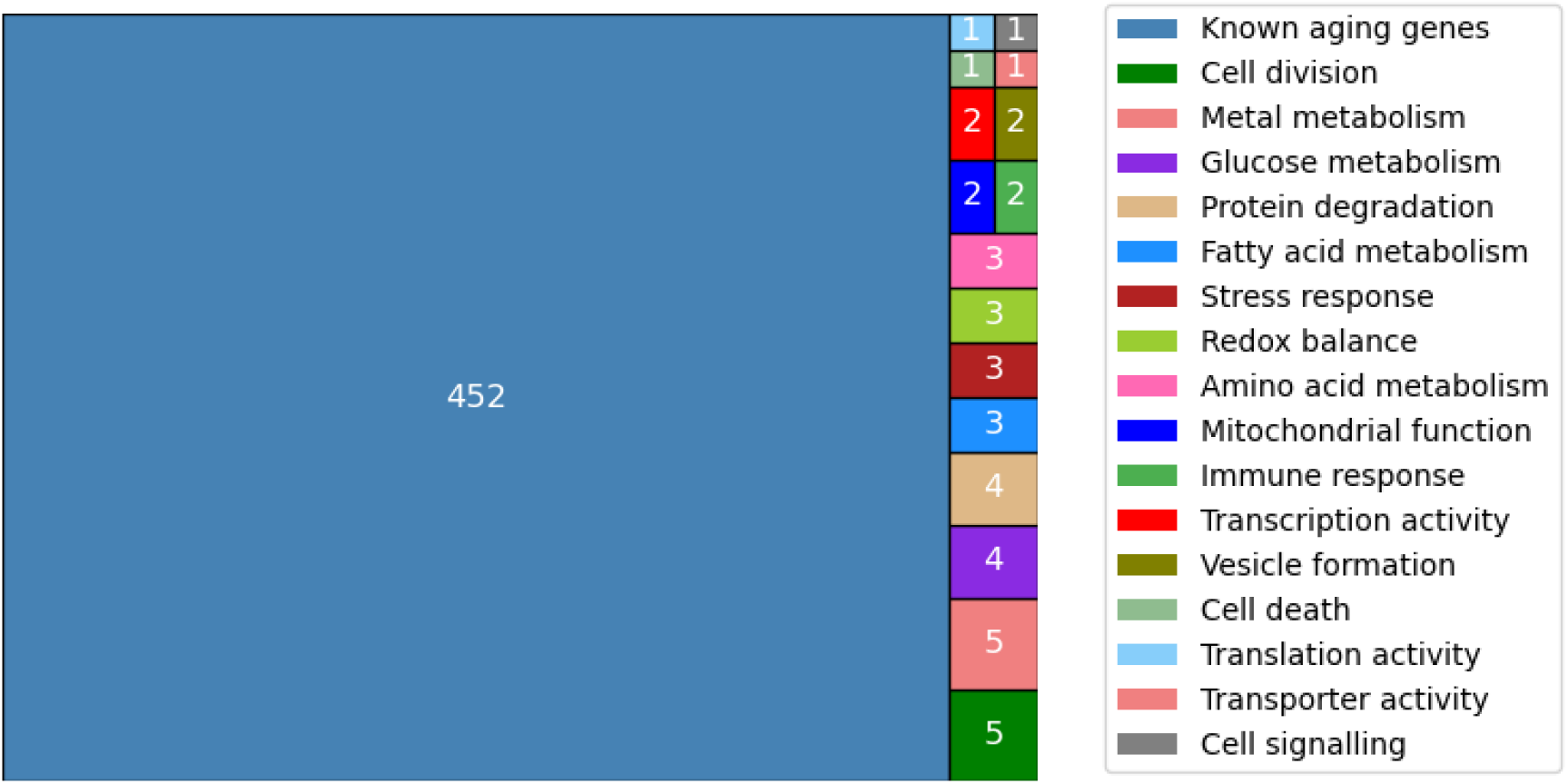
Summary of the top quantile of the ranked genes (Q1 in Figure 4). 91% (452) of the top ranked genes are known aging-related genes (6,34), while 9% (42) are not previously known to be associated with aging. The functional categories, based on Uniprot’s gene descriptions (35), of the potential new aging-related genes are listed.

### rsks-1 is similar to several dysfunction marker proxy genes

The gene rsks-1 shows an overall high ranking (top 2%) and is highly ranked with the three aging dysfunction types (morphological aging feature marker rank: 6%; intestinal aging marker rank: 6%; Aging-related gene rank: 1%) (Table 3). This high ranking is determined by the large degree of similarity of rsks-1 with several dysfunction marker genes across the different types of aging dysfunction markers (see Figure 6 for similarity with different types of aging dysfunction markers and Figure 7 for individual gene similarity). The aging-related genes showed the highest ranking score for rsks-1 (Figure 6), based on the phosphorylation data, biotype categorical data, gene ontology information, as well as the expression patterns in tissue types and different life stages.

**Figure 6:**
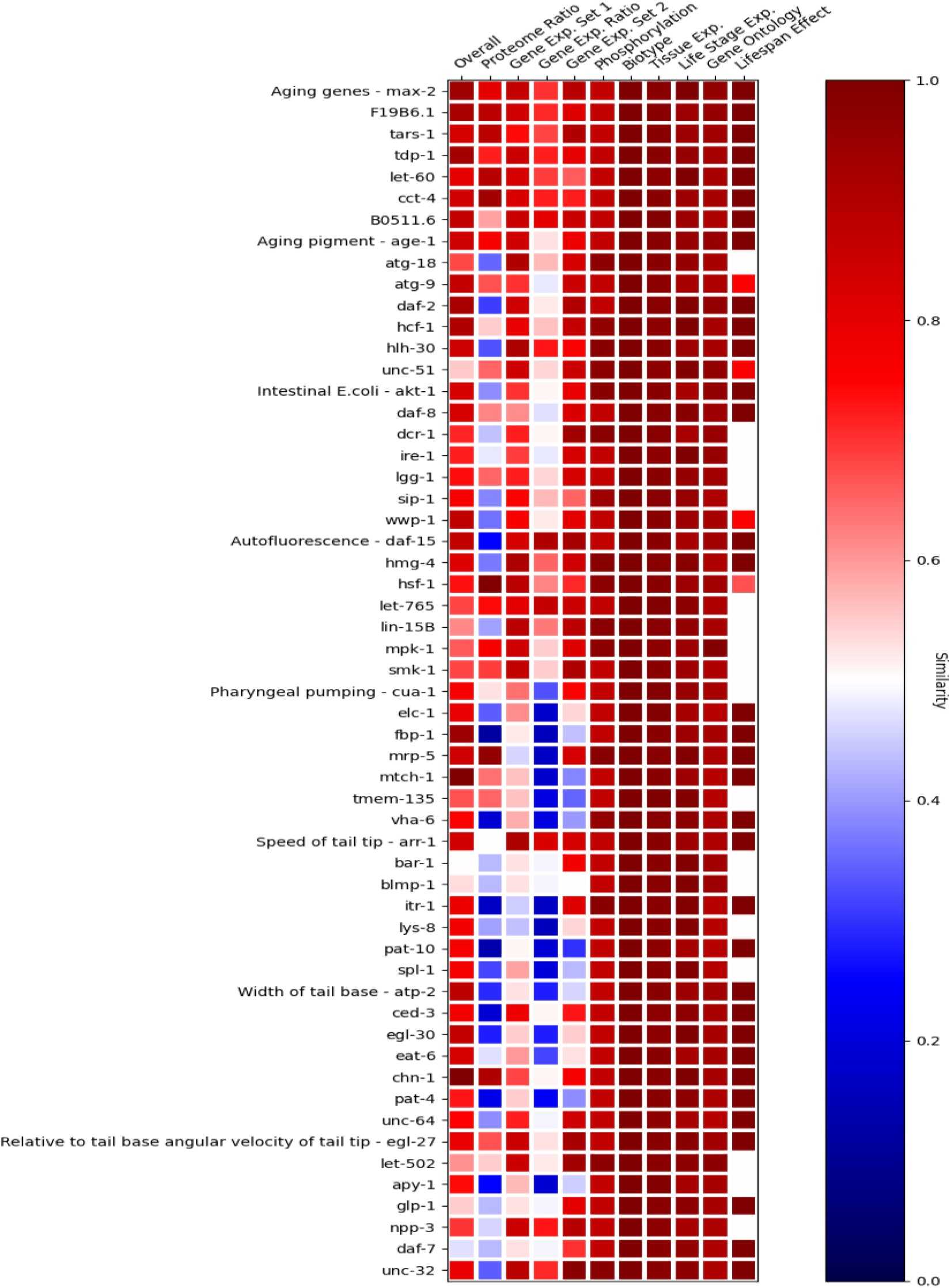
Correlation between rsks-1 and all dysfunction marker genes (rows) per datafield (columns 2 - 11), and overall similarity (first column) contribution to rsks-1’s ranking by each dysfunction marker gene.

**Figure 7:**
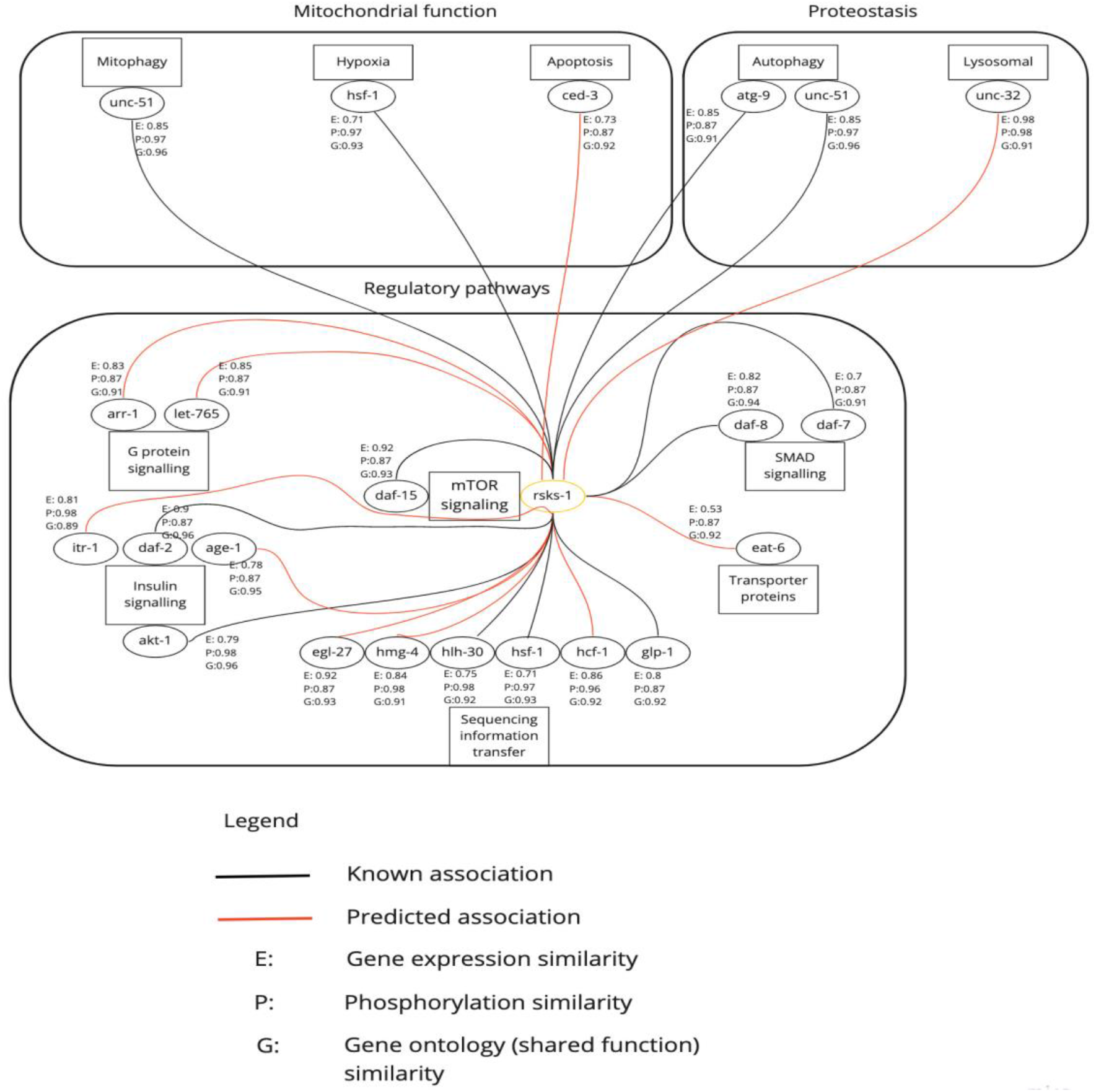
rsks-1’s known and predicted associations with the dysfunction marker genes based on the computationally determined similarity of gene expression (E), phosphorylation (P) and shared function from gene ontology (G). Black lines denote known associations based on literature, whereas red lines denote newly predicted associations.

### Dysfunction marker genes linked to rsks-1

In Figure 7, the known and predicted interactions between rsks-1 and similar dysfunction marker genes in their associated aging processes and systems based on UniProt keywords are displayed (35). Based on computational output, rsks-1 was highly similar to genes associated with mitochondrial function, redox balance, proteostasis and regulatory pathway systems. Based on existing literature and database searches (e.g. Pubmed, Wormbase, UniProt, https://www.uniprot.org/, and String, https://string-db.org/), rsks-1 and the following genes have a known interaction or are present in interacting pathways: unc-51, hsf-1, atg-9, daf-7, daf-8, glp-1, hlh-30, akt-1, daf-2, daf-15 (46–52). rsks-1 is predicted by the computational output to be highly associated with the following genes: ced-3, unc-32, eat-6, hcf-1, hmg-4, egl-27, age-1, itr-1, let-765, arr-1, with no current supporting literature of a direct association.

## Discussion

As our understanding of the aging process improves, the number of potential aging-related targets increases. This leads to the challenge of prioritizing targets of the aging process to promote successful aging and prevent the development of age-related diseases. Moreover, several functional parameters and the overall context of molecular targets in aging, is poorly reflected in current approaches to identify novel targets of interest. In this study, we propose a pipeline that integrates the plethora of publicly available genomic, transcriptomic, proteomic and morphological data of *C. elegans* with a supervised machine learning approach to prioritize aging-related genes. These ranked genes can be translated to known human orthologs potentially uncovering previously unknown information on the basic aging processes in humans. These genes could also serve as targets against aging-related diseases, such as AD. To test the capability of the pipeline, we used a known aging-related gene (rsks-1), enabling us to contextualize our findings with current literature.

### The computational output ranked aging-related genes of *C. elegans* highly

We repurposed recursive feature elimination with SVM and GLM models to develop a machine learning technique which ranks genes by their potential involvement in the process of aging. The computational output was validated by identifying the ranks of well-known aging-related genes such as age-1, daf-2, and let-363, all of which were ranked within the top 3% of the 4380 *C. elegans* genes. rsks-1, a ribosomal S6 kinase gene, was also ranked highly by our machine learning technique due to its high similarity with the identified proxy genes (associated with the dysfunction markers) and genes associated with aging processes. The overall ranking of rsks-1 was 55 out of 4380 genes (top 2% of all *C. elegans* genes), and it had the highest similarity with other genes associated with aging processes (n = 39, top 1%) (Table 3). The ranking of the known aging-related genes and rsks-1 is comparable to the GenAge ranking of longevity genes based on their ability to extend lifespan, however, rsks-1 is ranked much higher by our pipeline (6,34). This could be due to GenAge only ranking genes based on the percentage increase in lifespan after gene modulation, rather than the involvement of the gene in the aging process.

The *C. elegans* gene daf-2 (insulin-like receptor) is well-studied in the aging field and its knockout increases lifespan by 169%, whereas knockout of rsks-1 results in a lifespan increase of 20% (46). The dual knockout of daf-2 and rsks-1 has shown a synergistic lifespan extension by 454% (46). However, the mechanisms through which rsks-1 affects the aging process and results in this synergistic effect remains unclear. Characterizing the mechanisms involved in rsks-1 functioning will help discover and detail associated targets involved in the aging process, which could promote successful aging if inhibited or activated/stimulated.

Our pipeline identified other aging-related genes possibly related to rsks-1’s mechanism of aging. The genes from aging processes which were uncategorized (not directly associated with dysfunction markers) were used to detect similar aging-related genes, such as rsks-1. Additionally, proxy genes from the other dysfunction marker types (intestinal aging markers and morphological aging features) were used to predict possible associations (Figure 7) through which rsks-1 could affect the aging process in *C. elegans*.

### The potential interaction of rsks-1 with aging-related genes

The interactions between the proxy genes and rsks-1 with a similarity score were assessed using publicly available databases (Pubmed) (Figure 7). These genes include atk-1, daf-15, hlh-30, hsf-1, glp-1, daf-8, unc-51, atg-9, ced-3, unc-32, eat-6, hcf-1, hmg-4, egl-27, age-1, itr-1, let-765, arr-1.

There is supporting evidence for multiple of the associations proposed by our pipeline between the proxy genes and rsks-1 based on previous studies. Indeed, the String database (52) has predicted a functional association between rsks-1 and the dysfunction markers atk-1 and daf-15, based on their putative homologs interacting in other organisms. A possible relationship has been shown between hlh-30 and rsks-1 in *C. elegans*, with knockout of rsks-1 resulting in increased hlh-30 mRNA levels compared to wild type (51). Further, a specific genetic interaction has been found between rsks-1 and hsf-1 through RNA interference screening (50). Through an RNA interference study it was shown that unc-51 and atg-9 extended the lifespan of *C. elegans* with an rsks-1 mutant (47). There is also a functional interaction between rsks-1 and the dysfunction marker glp-1, in which rsks-1 promotes glp-1 fate; the nature however, of this interaction, remains unclear (49). A null mutation with rsks-1 and daf-8 showed a strong synergistic Daf-c phenotype (48), further supporting the association between the proxy genes and rsks-1 using the similarity score from our pipeline.

The remaining dysfunction marker genes were to the best of our knowledge not previously described in literature to have an interaction or association with rsks-1. These include the genes: ced-3, unc-32, eat-6, hcf-1, hmg-4, egl-27, age-1, itr-1, let-765, and arr-1. Our computational output indicates a possible interaction or association between the latter genes and rsks-1 based on a combination of coexpression, phosphorylation pattern, and functional description (categorical data). The potential relationship between the proxy genes and rsks-1 could allude to new mechanisms through which rsks-1 affects aging in *C. elegans*. Our pipeline, therefore, does not only rank aging-related genes, but also provides valuable information on how the ranked genes might interact or be associated with the proxy genes based on their similarity score.

### Using cellular and morphological dysfunction to describe effects of targets on the aging process and possible relevance to human disease

When implementing the pipeline presented here, the proxy genes related to dysfunction markers enable the detection of aging-related genes that have a large potential of affecting aging outcomes. This is due to the close association between their gene expression and the change in the dysfunction marker over the lifespan of *C. elegans*. Although gene expression is not indicative of protein activity, for our purposes it is an initial indicator of a possible relationship between a gene and an aging-related feature (**Supporting Figures**: see Supplementary information).

As an example, daf-2 was selected as a proxy gene for aging pigment due to its high RMSE value when comparing its expression with aging pigment accumulation. Knockout of daf-2 has resulted in a decreased accumulation of aging pigment and improved locomotory function (22), supporting its relevance to the presence of aging pigment. Similarly, in our computational output rsks-1 shows the highest similarity to genes related to aging pigment and tail tip movement. Both daf-2 and rsks-1 impact *C. elegans* lifespan through shared pathways, ultimately involving translation regulation (53). Therefore, it is expected that they would have similar effects on molecular and morphological aging features, which is reflected in the computational output.

Aging pigment forms due to a combination of dysregulation in proteostasis, as well as disturbances in redox balance. Locomotion (such as tail tip movement) is related to aging pigment formation (22). These functional outcomes could be equivalent to dysregulation of cellular function in neuronal cells, such as amyloid beta aggregation in humans. As changes in rsks-1’s function may lead to improvement (or deterioration) in the healthspan of *C. elegans* during aging, this may suggest that targeting its human homolog S6 kinase in neurons could prevent the accumulation of proteins that contribute to the onset of neurodegenerative diseases. This is supported by the observation that a genetic reduction in S6K1 reduced the generation of amyloid beta in mice (54). Therefore, the pipeline presented here is able to detect and describe potential aging-related genes relevant to AD.

## Conclusion and future outlook

The aim of this study was to develop a computational pipeline enabling to prioritize *C. elegans* genes by their probability of being involved in aging-related functional processes. This was achieved by using a supervised machine learning technique to rank genes by their similarity to known aging-related genes and dysfunction marker genes (known to be present in aging *C. elegans*). The ranked gene output showed that 91% of the top-ranked quantile of genes are known aging-related genes, while the remainder could be potential novel aging-related genes. The accuracy of the ranked output was shown through the high ranking of known aging-related genes, age-1, daf-2, let-363, and rsks-1.

Next, rsks-1 was used as an example gene to showcase the output’s functionality. The dysfunction marker (proxy) genes with a high similarity to rsks-1 could potentially indicate a functional interaction or association. Furthermore, the dysfunction markers (eg. aging pigment, tail tip speed, etc.) could be used to understand how a highly ranked gene may affect the aging process. Throughout, the computational output was validated through contextualisation with the most recent literature.

Future work that includes information beyond gene expression may further strengthen approaches similar to the pipeline presented here, to describe the connection between genes and dysfunction markers (such as protein and phosphorylation data). In addition, causal inference analysis, as described in the works of Pearl *et al*., 2016, 2019, may add value by uncovering the causal structure in the system, allowing one to identify potential new treatment targets not evident from correlation and machine learning analyses (55,56).

Well-designed experiments that are guided by the current computational output may inform current gaps, such as the association between proxy genes and dysfunction markers, which will improve the relevance of the identified age-related targets. Such approaches may allow a more in-depth characterization of the association between gene expression and dysfunction marker decline. Finally, by using a workflow similar to that demonstrated in the present study for rsks-1, unknown genes could be explored and prioritized to potentially identify novel targets for aging.

## Acknowledgements

We would like to acknowledge Prof Ben Loos and Dr. Dawie van Niekerk for assistance with formatting and editing of the manuscript, and Dr. Anthony Sedgwick for assistance with the initial conceptualization of the study.

## Contributions

NT and ZJVR: Data collection, analysis, and interpretation of results. CL: Project design and development, computational analysis. RO provided computational and mathematical advice. RS and all authors contributed to manuscript writing and revision. All authors read and approved the final manuscript.

