## Supplementary information for "Supervised machine learning with feature selection for prioritization of targets related to time-based cellular dysfunction in aging"

A

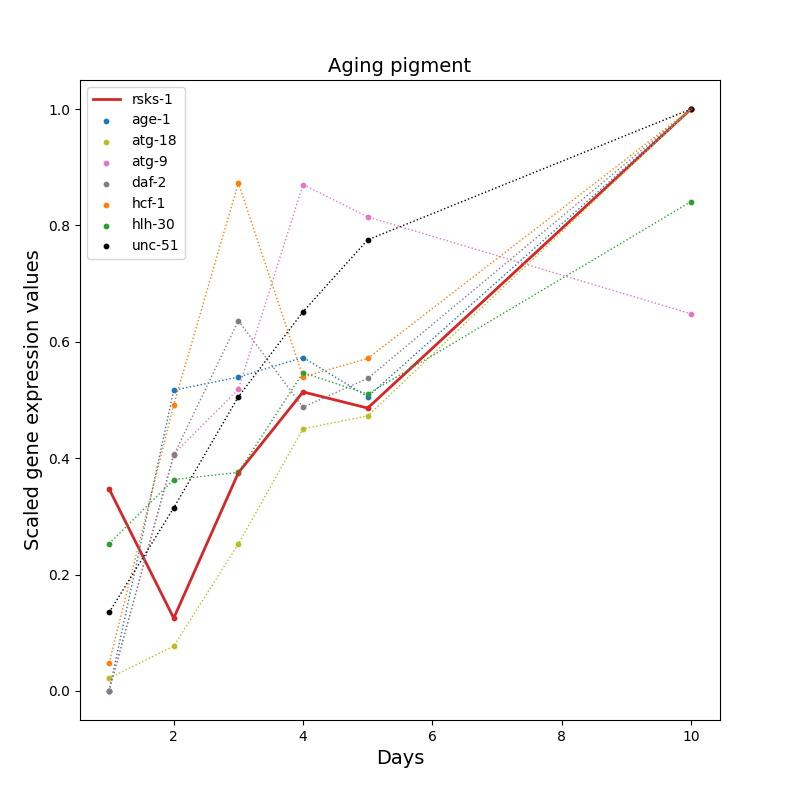

C
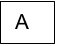

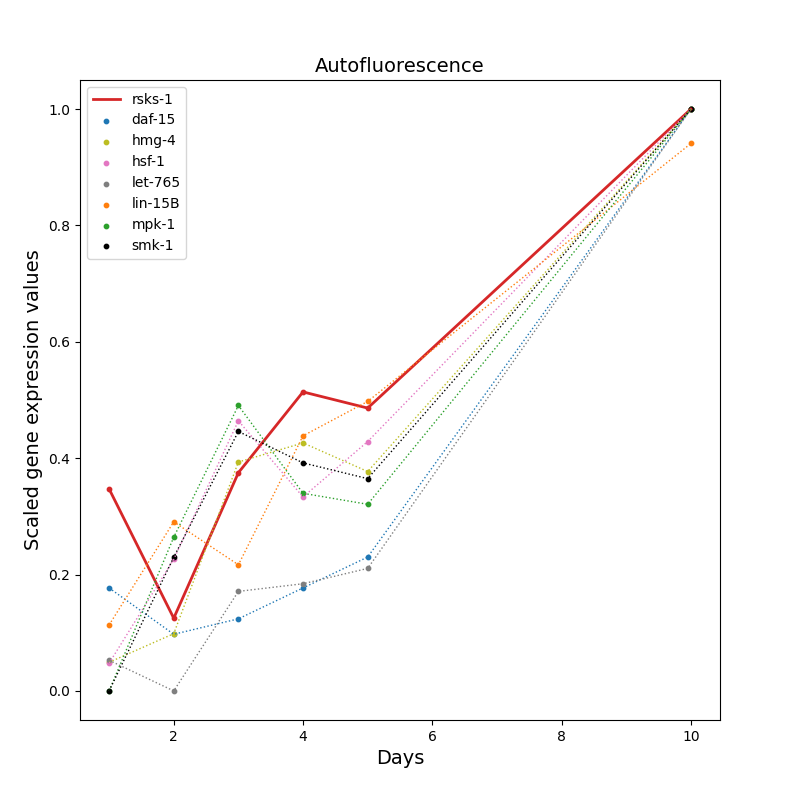

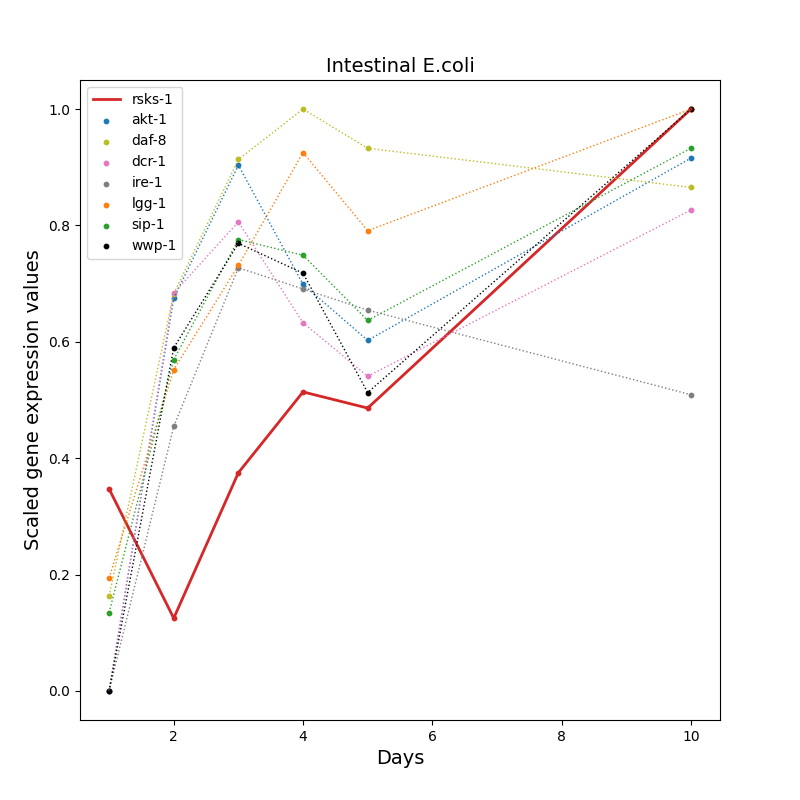

B
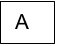

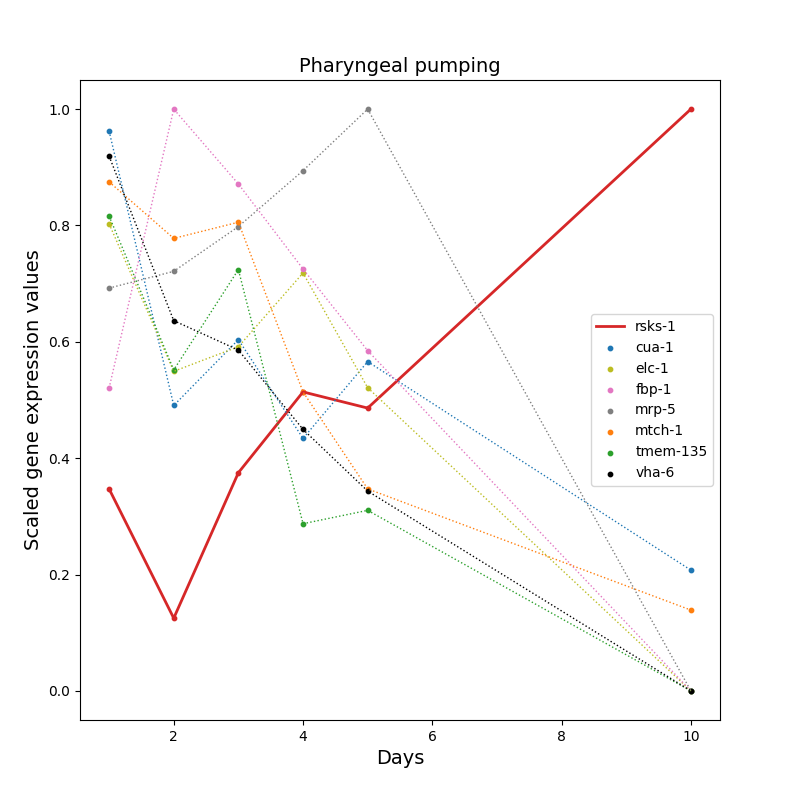

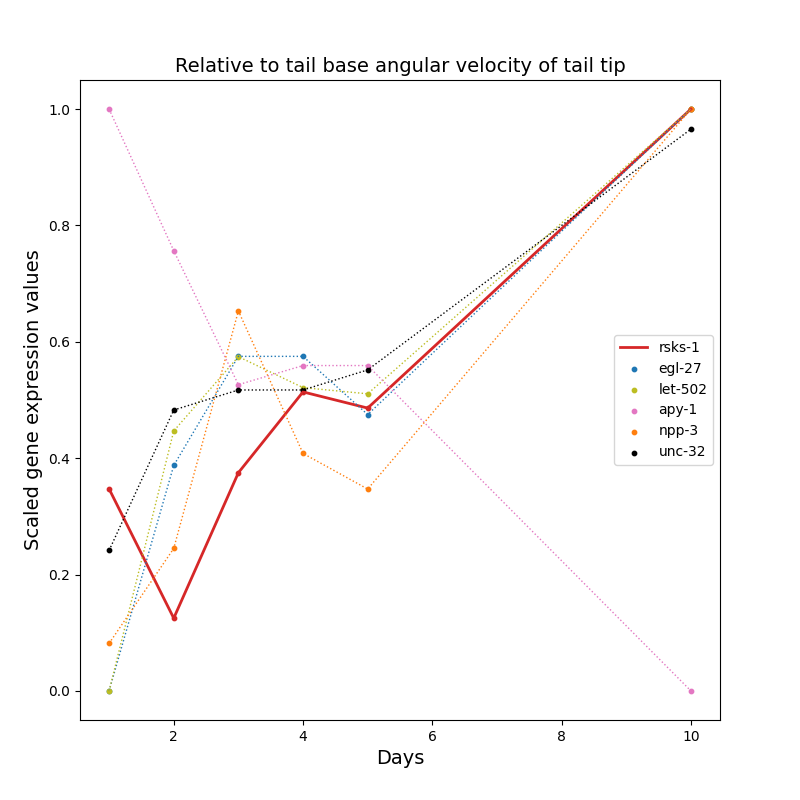

G
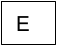

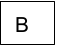

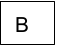

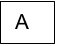

F
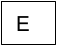

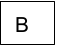

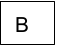

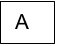

E
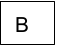

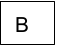

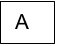

D
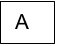
v

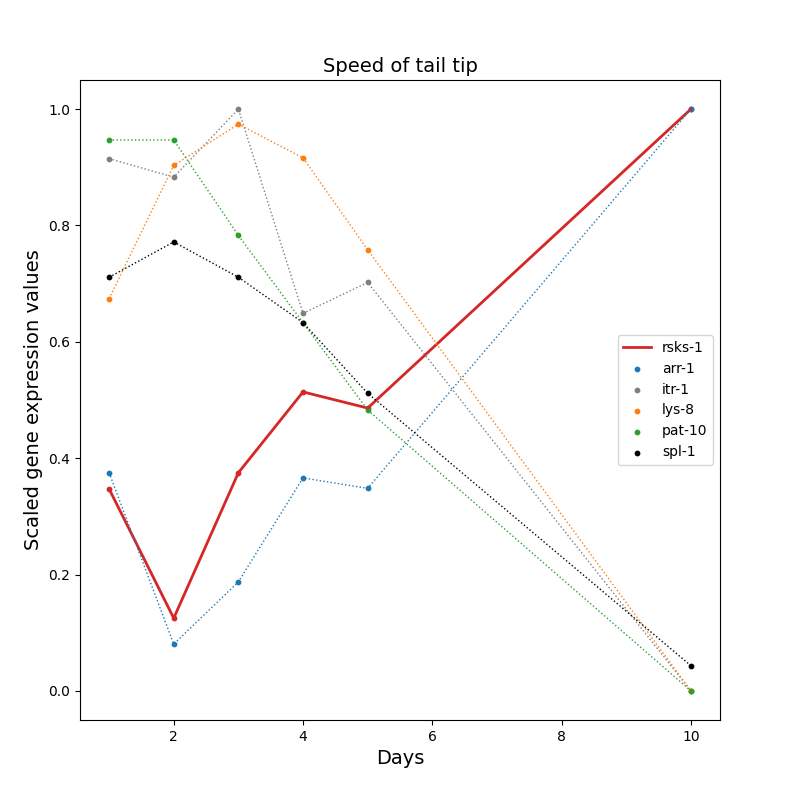

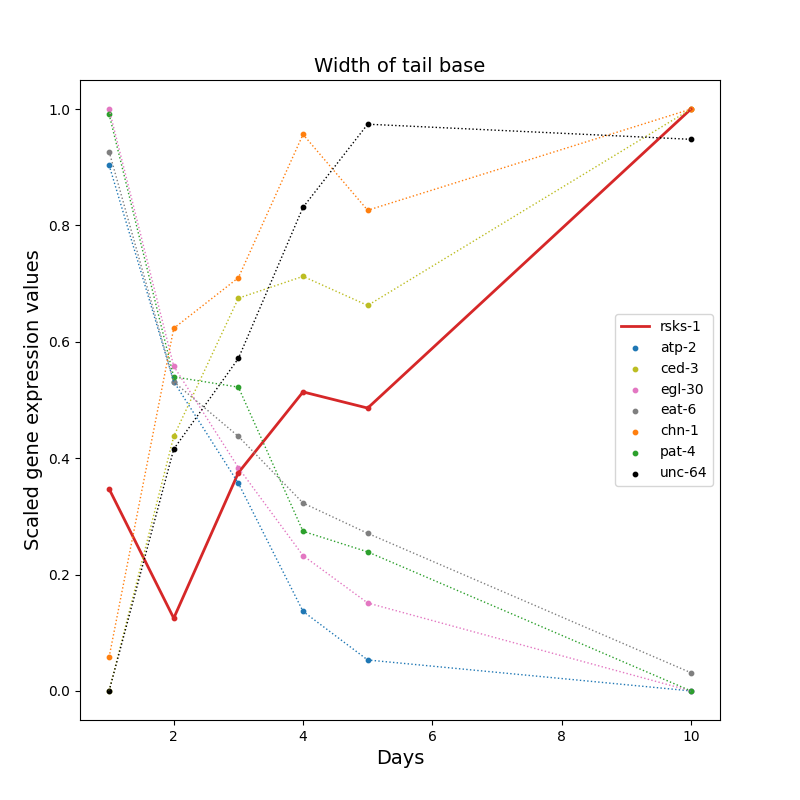

*Supplementary Figures: Change in gene expression dataset 1 over time for rsks-1 (red) and dysfunction marker proxy gene for: (A) Aging pigment (B) Autofluorescence (C) Intestinal E. coli (D) Pharyngeal pumping (E) Relative to tail base angular velocity of tail tip (F) Speed of tail tip (G) Width of tail base.*
